# Characterization of sperm motility signaling pathway in a gonochoric coral suggests conservation across sexual systems

**DOI:** 10.1101/2022.12.15.520596

**Authors:** Benjamin H. Glass, Jill Ashey, Amarachukwu R. Okongwu, Hollie M. Putnam, Katie L. Barott

## Abstract

Many marine invertebrates liberate their gametes into the water column broadcast spawning, where fertilization hinges upon the successful activation of sperm motility. Here, we investigated the molecular mechanisms underpinning sperm motility in the broadcast spawning coral *Astrangia poculata*. We found that cytosolic alkalinization activates the pH-sensing enzyme soluble adenylyl cyclase (sAC), followed by motility, in *A. poculata* sperm. In addition, we show for the first time in any cnidarian that sAC activity is required to activate protein kinase A (PKA) in sperm, and that PKA activity is required for the initiation of sperm motility. Ultrastructures of *A. poculata* sperm displayed morphological homology to other gonochoric cnidarians, and investigation of cnidarian proteomes revealed that sAC, the central signaling node in the sperm motility pathway, demonstrates broad structural and functional conservation across a diversity of cnidarian species. Ultimately, these results suggest that the role of sAC signaling in sperm motility is conserved between sperm from gonochoric and hermaphroditic corals, which is surprising given their morphological dissimilarities. This study also offers insight into the evolution of the mechanisms controlling metazoan sperm motility.

**Summary statement:** For broadcast spawning marine invertebrates, the initiation of sperm motility is essential for fertilization. Here, we provide evidence for conservation of a sperm motility pathway across sexual systems in corals.

## Introduction

Sexual reproduction is ubiquitous across the tree of life, and species have developed diverse mechanisms for the allocation of sexual cell types among individuals (i.e., sexual systems), such as gonochorism and hermaphroditism (Leonard, 2018). The phylum Cnidaria is comprised of a diverse assemblage of species (e.g. corals, sea anemones, and jellyfish) whose sexual systems are not always correlated with their phylogenetic relationships (Siebert and Juliano, 2017). For example, there are both gonochoric and hermaphroditic species within the cnidarian orders Scleractinia (stony corals) and Actiniaria (sea anemones), and even confamilial species can display different sexual systems (Harrison, 2011; Kerr et al., 2011; Reuven et al., 2021). Cnidarians also display variation in reproductive strategies, with some species exhibiting internal fertilization and brooding of larvae within the maternal polyp while others liberate both sperm and eggs into the water column, where sperm must perform taxis towards a conspecific egg in order to achieve fertilization (Kerr et al., 2011; Morita et al., 2010). While the sexual systems and reproductive strategies of many cnidarian species have been described (Baird et al., 2009; Harrison, 2011), we lack a thorough understanding of cnidarian sperm biology, which limits conservation strategies to protect these ecologically important species from the effects of anthropogenic climate change (van Woesik et al., 2022).

Early work investigating sperm in cnidarians revealed a wide diversity of morphologies presented by species in this phylum (Fadlallah, 1983; Gaino and Scoccia, 2010; Gaino et al., 2013; Goffredo et al., 2000; Hinsch, 1974; Hinsch and Clark, 1973; Padilla-Gamiño et al., 2011; Steiner, 1991; Steiner, 1993; Steiner and Cortés, 1996; Szmant-Froelich et al., 1980). This foundational work showed that cnidarian sperm could be categorized into distinct morphological “types,” which were found to be tightly associated with sexual system, in addition to differing within and between phylogenetic groups (Hinsch, 1974; Hinsch and Clark, 1973; Steiner, 1991). Specifically, sperm from hermaphroditic cnidarians tend to display rounded nuclei, whereas sperm from gonochoric species typically have conically shaped nuclei (Steiner, 1993). Additionally, sperm from gonochoric cnidarians show partial fusion of mitochondria, the presence of dense perinuclear vesicles, and a dense lipid body, whereas these features are absent in sperm from hermaphroditic species (Steiner, 1993). As sperm morphology is associated with function across phyla (Darszon et al., 2020), these differences in morphology are likely associated with different functional constraints between gonochoric and hermaphroditic cnidarian species. Indeed, sperm from hermaphroditic corals appear to have receptors for egg-derived immobilizing factors (in addition to chemoattractants), which have never been described in gonochoric species (Kaupp et al., 2006; Morita et al., 2006; Morita et al., 2009b; Yoshida et al., 2008), indicating that sperm molecular phenotypes may differ between coral sexual systems. Yet, it remains unknown whether sperm from gonochoric and hermaphroditic corals differ in terms of the molecular mechanisms that regulate their motility, a central function necessary for successful reproduction.

Recent work has shown that components of a molecular sperm motility signaling pathway involving the enzymes soluble adenylyl cyclase (sAC) and protein kinase A (PKA) is conserved between sea urchins and the hermaphroditic coral *Montipora capitata* (Speer et al., 2021). This “sAC-cAMP-PKA” signaling pathway has been most thoroughly described in sea urchin sperm, where the signaling cascade begins with the binding of an egg-derived substance (e.g. speract) to guanylyl cyclase A (GC-A; aka speract receptor), a sperm transmembrane receptor (Coll et al., 1994; Speer et al., 2021; Zhang et al., 2019). The activation of GC-A results in the production of the second messenger cyclic guanosine monophosphate (cGMP), which activates the potassium-selective, cyclic nucleotide gated channel (CNGK), resulting in the removal of H^+^ ions from the sperm cytosol (Kaupp et al., 2006; Morisawa and Yoshida, 2005). This leads to an increase in sperm internal pH (pH_i_) which activates the pH-sensing enzyme sAC (Nishigaki et al., 2014). Activated sAC then converts adenosine triphosphate (ATP) to cyclic adenosine monophosphate (cAMP), a second messenger which modulates the activity of PKA and other downstream targets to produce sperm motility (Nishigaki et al., 2014). Homologs of each component of the sea urchin pathway are expressed in sperm from the coral *M. capitata*, and sperm cytosolic alkalinization leads to the activation of adenylyl cyclases (ACs) and PKA, followed by the onset of sperm motility in this species (Speer et al., 2021). However, several aspects of this pathway remain undescribed in cnidarians. For example, it is still unknown whether AC activity during the onset of motility is due to sAC (as opposed to transmembrane ACs), whether sAC activity is necessary for activating PKA, or if PKA itself is necessary for sperm motility. It is also unknown whether some or all components of this pathway are conserved in gonochoric corals, which may be unlikely given the numerous morphological and molecular differences between sperm from gonochoric and hermaphroditic species. Here, we address this knowledge gap by employing a variety of molecular techniques to characterize the role of sAC and PKA in regulating sperm motility in the gonochoric coral *Astrangia poculata*. We then performed an analysis of published proteomes to investigate whether sAC displays broad structural and functional conservation across cnidarian species with diverse sexual systems.

## Results

### Sperm ultrastructural characterization via TEM

Transmission electron microscopy (TEM) was used to characterize the ultrastructure of *Astrangia poculata* sperm collected from spawning male colonies (N = 3 males; Fig. 1A–C). TEM micrographs showed that sperm had a head size of approximately 2.5 μm (greatest width) by 3 μm (greatest length), and displayed a tail length of approximately 45 μm (Fig. 2A). At the base of the sperm head, 2–4 mitochondria formed a semicircular structure around the centriolar complex, which gave way to the flagellar axoneme at the base of the head (Fig. 2B–F). An electron dense lipid body was located planar with the mitochondria; together, these structures formed a full ring around the base of the sperm head, which filled the cytoplasmic space from the outer plasma membrane to the small centriolar complex (Fig. 2D, F). A transverse section of the flagellar axoneme revealed microtubules in the classical 9+2 arrangement, and pericentriolar processes could be faintly seen connecting the central microtubule pair to the outer microtubule doublets (Fig. 2E). The nucleus was conically shaped, located anterior to the mitochondria and lipid body, and included a region of low electron density at the anterior tip (Fig. 2B, F). The tip of the nucleus was directly adjacent to the outer plasma membrane. In contrast, more cytoplasmic space existed between the nucleus and plasma membrane near the nuclear base (Fig. 2B, F). This space was mostly devoid of visible structures, but some electron dense vesicles were observed in this region (Fig. 2F), as well as in the space between the mitochondria and the outer membrane (Fig. 2B–D, F).

**Figure 1.**
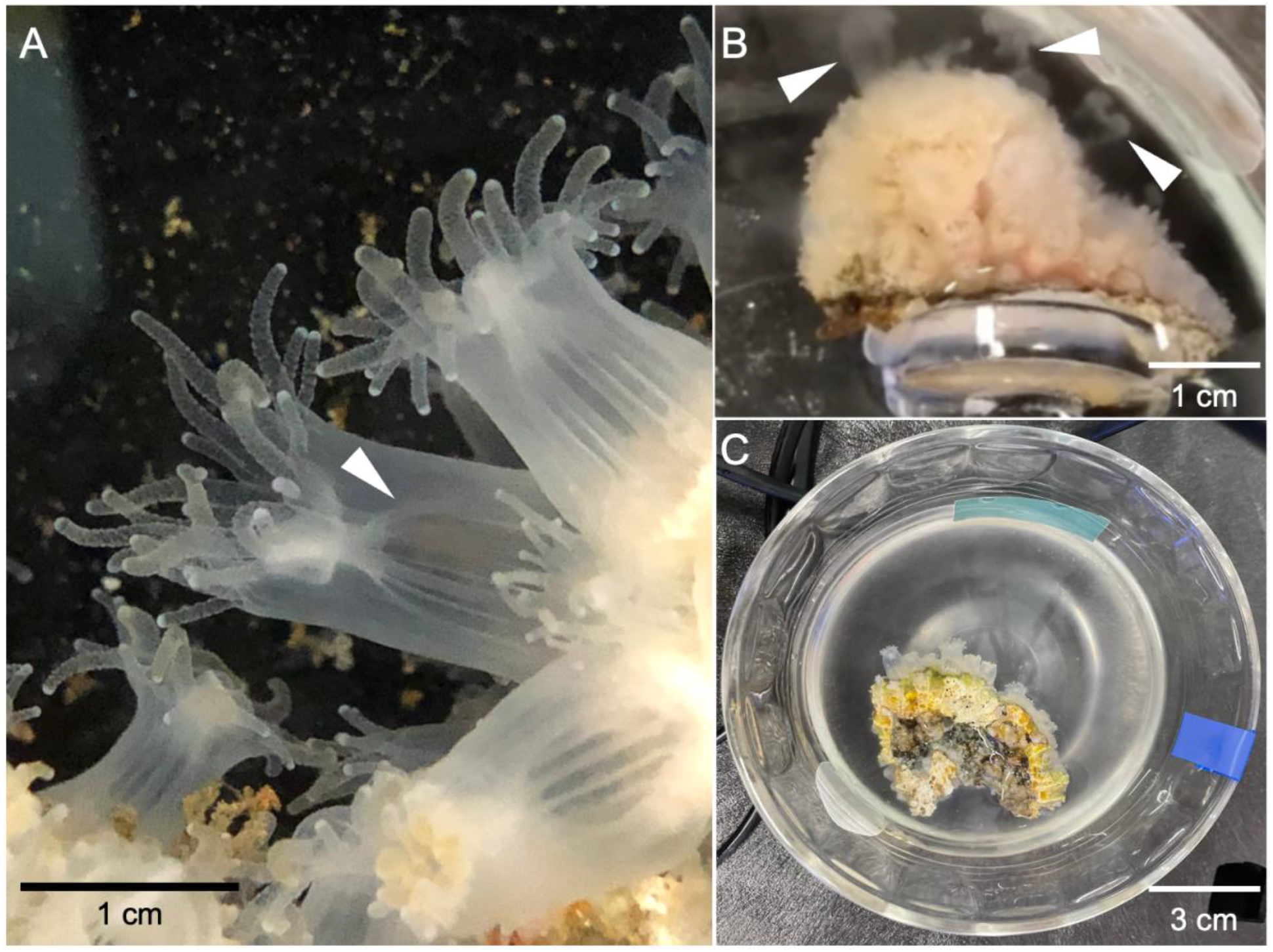
*Astrangia poculata* colony and spawning images. A) A close-up image of several polyps of *A. poculata* in its aposymbiotic form. White arrowhead indicates one polyp’s mesenteries (the site of gametogenesis). B) An aposymbiotic, male colony of *A. poculata* actively spawning sperm, which are seen being released from the polyps in concentrated bursts (white arrowheads). C) An aposymbiotic, male colony of *A. poculata* sitting in sea water cloudy with recently spawned sperm. Scale bars are included in all images.

**Figure 2.**
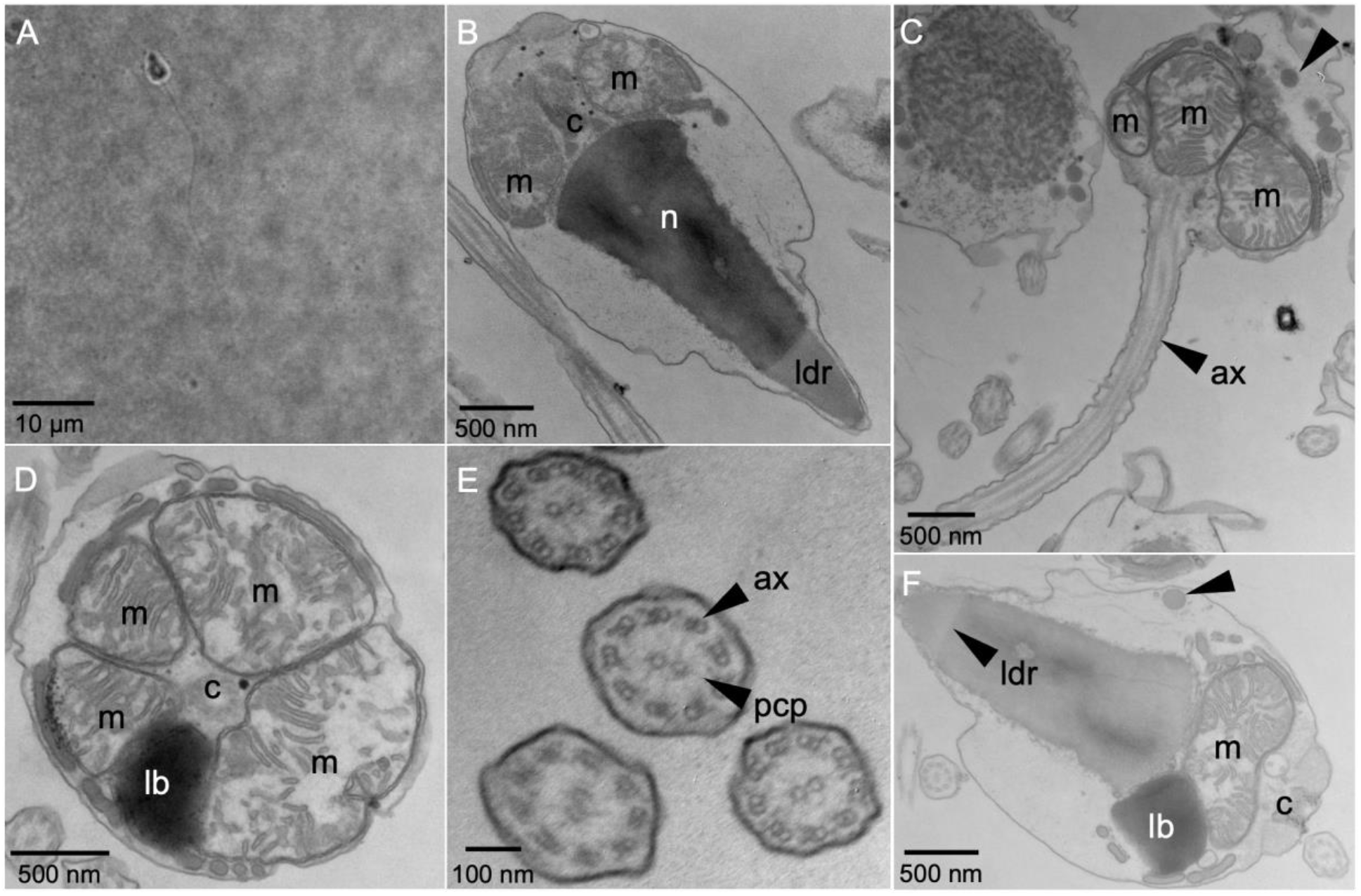
*Astrangia poculata* sperm ultrastructure. A) A brightfield image of a single sperm cell. (B-F) Transmission electron micrographs of *A. poculata* sperm. B) A sagittal view of the internal structure of a single sperm head. Labeled structures include the region of low density (ldr) at the anterior tip of the nucleus (n), two mitochondria (m), and the centriolar complex (c). C) A parasagittal view of a sperm cell depicting the base of the head leading into the tail. Labeled structures include the flagellar axoneme (ax), three mitochondria (m), and perinuclear vesicles (e.g. black arrowhead). D) A transverse view of the base of a sperm head depicting four mitochondria (m), a dense lipid body (lb), and the centriolar complex (c). E) A transverse view of several sperm flagella depicting axonemes (ax) with their classical 9+2 microtubule arrangement. Pericentriolar processes (pcp) are somewhat visible between the central microtubule pairs and the outer microtubule doublets of each flagellum. F) A sagittal view of the internal structure of a sperm head, again depicting the anterior region of lower density (ldr), dense lipid body (lb), mitochondrion (m), centriolar complex (c), and vesicles (black arrowhead). Scale bars are included in all images.

### Expression, localization, and activity of sAC in sperm

We set out to characterize aspects of the sea urchin sperm motility pathway (Fig. 3A) in *A. poculata* sperm. Western blotting for soluble adenylyl cyclase (sAC) in *A. poculata* sperm revealed expression of a single major isoform approximately 88 kDa in weight (N = 11 males; Fig. 3B). Immunocytofluorescence revealed that sAC was localized mostly at the base of the sperm head, though it was also present in low density throughout the sperm tail (N = 4 males; Fig. 3C–D; Fig. S1A–B). Baseline levels of cAMP were 25 ± 4 nmol cAMP per ng protein in DMSO treated sperm (Fig. 3E). Addition of 20 mM NH_4_Cl (which induces cytosolic alkalinization) resulted in a 3-fold increase in cAMP content within six seconds, which peaked at 75 ± 14 nmol cAMP per ng protein. This level began to decrease by 30 seconds post-stimulation, returning to just above the baseline at 33 ± 7 nmol cAMP per ng protein by five minutes (Fig. 3E). In contrast, in the presence of the sAC inhibitor KH7, there was no increase in cAMP concentration at six seconds following stimulation with 20 mM NH_4_Cl, and cAMP levels remained near the baseline at five minutes post-stimulation (Fig. 3E). Stimulation with NH_4_Cl (Type III ANOVA; *p* < 0.001), time (*p* < 0.001), and the interaction between treatment (DMSO or KH7) and stimulation (*p* < 0.05) were significant predictors of sperm cAMP levels (N = 9 assays per treatment).

**Figure 3.**
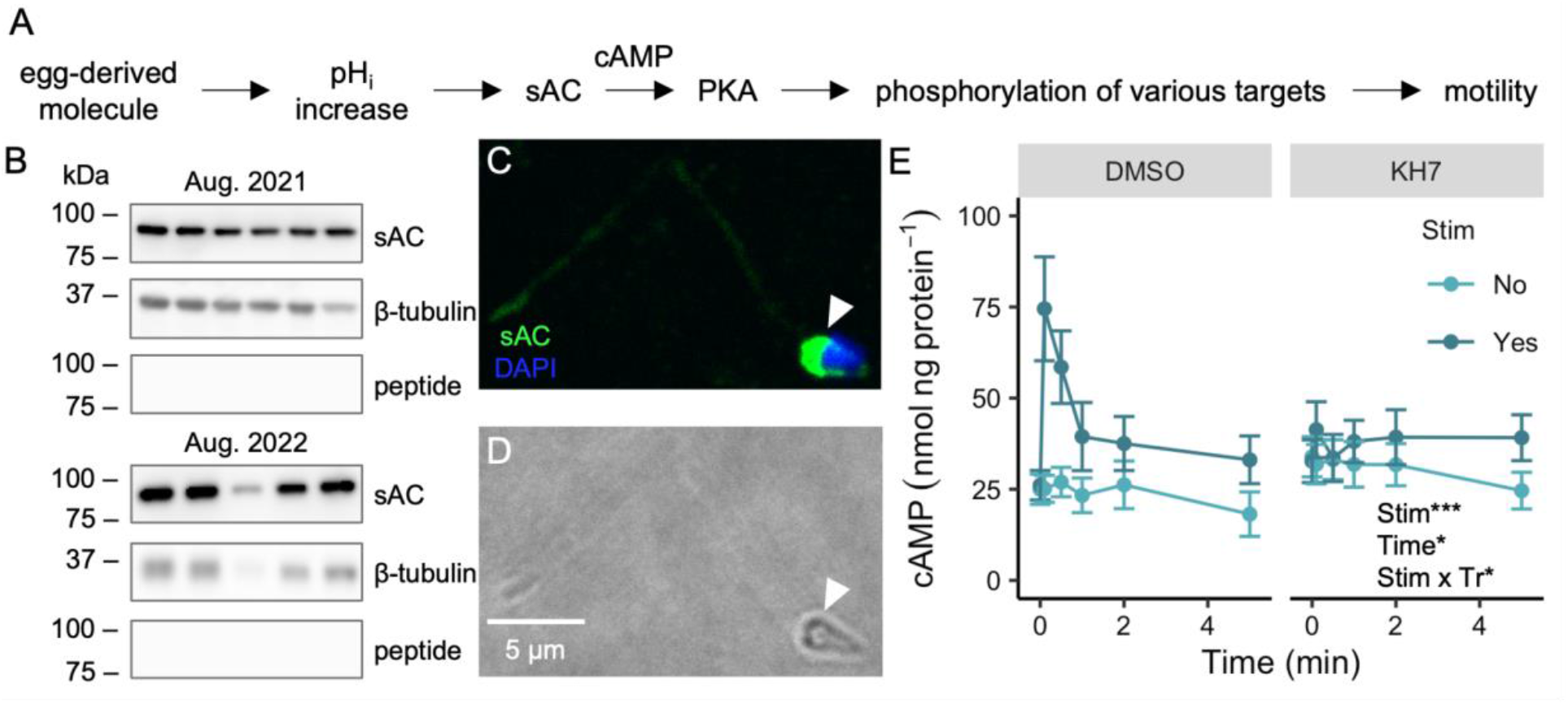
Sperm motility pathway and expression, localization, and activation dynamics of soluble adenylyl cyclase (sAC) in *Astrangia poculata* sperm. A) Simplified schematic of the sea urchin sperm motility pathway. B) Western blots for sAC, β-tubulin (loading control), and sAC again following antibody-peptide pre-absorption (nonspecific binding control) in sperm samples across two spawning seasons (Aug. 2021 and 2022). Each lane represents sperm from a distinct male colony (N = 11 males). C) Representative immunocytofluorescence image of a single sperm cell. Labels indicate staining for sAC (green) and DNA (DAPI; blue). White arrowhead (right) highlights sAC staining at the base of the sperm head. D) Brightfield image corresponding to image in panel B; scale bar applies to both images. E) Sperm *in vivo* cAMP concentration over time following incubation with DMSO or the sAC inhibitor KH7 and in the presence or absence of 20 mM NH_4_Cl stimulation (“stim”), which was administered immediately after time = 0 min. Each point represents the average of cAMP concentrations for pools of sperm from three distinct male colonies, which were each assessed in triplicate as technical replicates for a total of N = 9 assays per point (N = 216 assays in total). Bars represent the standard error of the mean, and the inset identifies stim, time, and the interaction between stim and drug treatment (Tr) as significant linear model terms (Type III ANOVA; * = *p* < 0.05, *** = p < 0.001).

### Structure, expression, and activity of PKA in sperm

We investigated the proteome of *A. poculata* and located two isoforms of the Cα subunit of protein kinase A (PKA), which differed in a few C-terminal amino acids and were estimated to be 40.28 and 40.25 kDa in weight. Generation of 3D structures for both PKA Cα isoforms by AlphaFold (Jumper et al., 2021) showed that the two proteins were nearly identical in structure, with the lighter isoform lacking a small helix at the C-terminus compared to the heavier isoform (Fig. 4A). Western blots showed that *A. poculata* sperm expressed both isoforms of PKA Cα, with the lighter isoform displaying slightly higher expression compared to its heavier counterpart (N = 6 males; Fig. 4B). In sperm treated with DMSO as a carrier control, stimulation with 20 mM NH_4_Cl resulted in increased PKA activity within six seconds, as evidenced by an increased number and darkening of bands on a Western blot treated with antibodies detecting phosphorylated PKA substrates (Fig. 4C). Quantification of the cumulative Western blot band intensity indicated that PKA activity increased rapidly following stimulation with 20 mM NH_4_Cl, peaking at 1.47 ± 0.04 times baseline levels by 30 seconds (Fig. 4D; Fig. S1C–D). Furthermore, levels of PKA substrate phosphorylation remained elevated at around 1.14 ± 0.05 times baseline levels for at least five minutes post-stimulation for sperm treated with DMSO (Fig. 4D). Incubation of sperm with the sAC inhibitor KH7 prior to stimulation changed the kinetics and intensity of PKA activation. Specifically, sperm treated with KH7 showed a 1.2-fold smaller peak in PKA substrate phosphorylation relative to controls of only 1.25 ± 0.02 times baseline levels, which was observed at six seconds following stimulation, as opposed to the peak at 30 seconds observed in sperm treated with DMSO (Fig. 4D). Furthermore, levels of PKA substrate phosphorylation decreased back to baseline (1.00 ± 0.04) by two minutes following stimulation in KH7 treated sperm, and were below baseline levels (0.92 ± 0.03) at five minutes (Fig. 4D). When treated with the PKA inhibitor H-89 prior to stimulation, sperm displayed 1.3-fold lower PKA substrate phosphorylation activity relative to DMSO controls, with a peak at 1.16 ± 0.02 times baseline levels occurring at 30 seconds before a return to just below baseline (0.96 ± 0.03) by five minutes (Fig. 4D). Time (Type III ANOVA; *p* < 0.05) and the interaction between time and treatment (*p* < 0.001) displayed a statistically significant relationship with PKA substrate phosphorylation (N = 18 assays per drug treatment).

**Figure 4.**
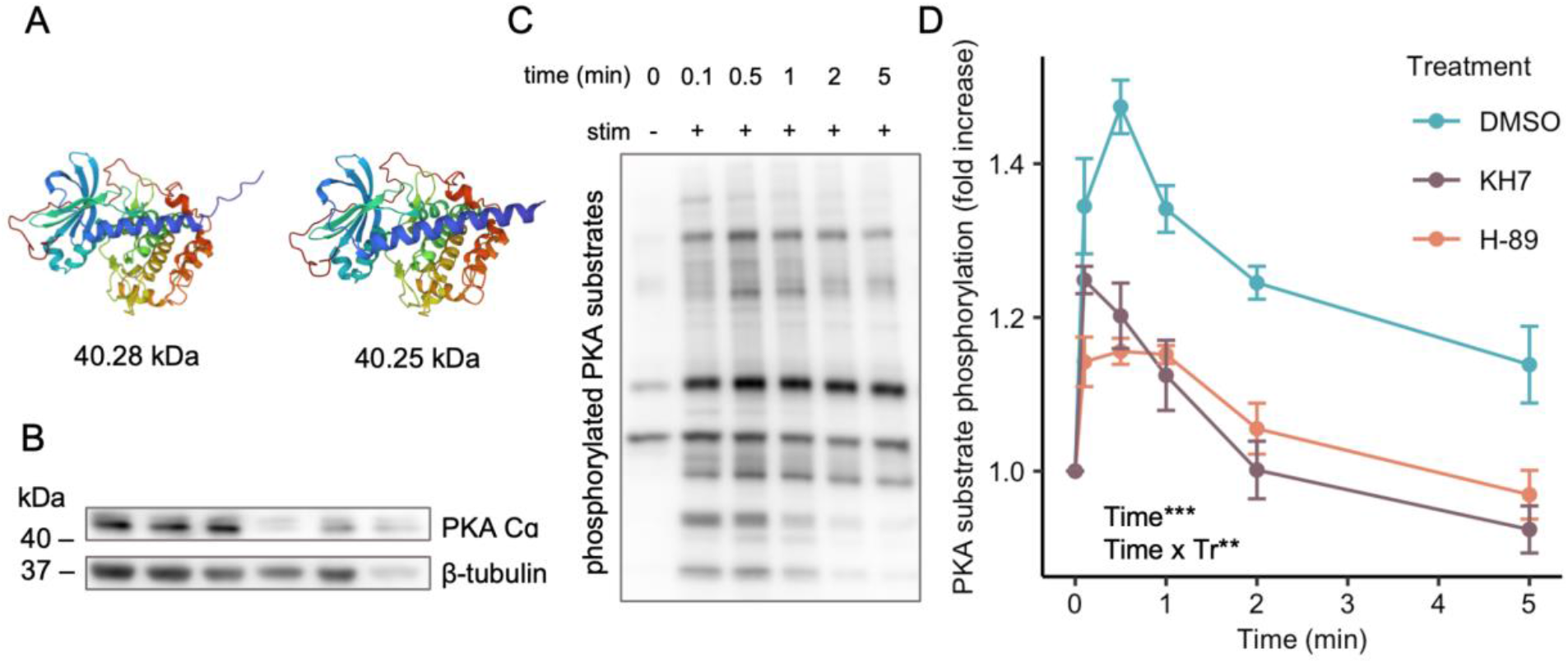
Structure, expression, and activation dynamics of protein kinase A (PKA) Cα in *Astrangia poculata* sperm. A) Predicted 3D structures for two isoforms of PKA Cα identified from the *A. poculata* proteome, with their corresponding predicted molecular weights. B) Western blot for PKA Cα displaying two bands just above 40 kDa, which are assumed to be the isoforms of PKA Cα depicted in panel A. Blot for β-tubulin is also shown as a loading control. Each lane represents sperm from a unique male genotype (N = 6 males). C) Representative Western blot for phosphorylated PKA substrates over time following sperm stimulation with 20 mM NH_4_Cl (“stim”), which was administered directly after time = 0 min. A pool of sperm from a single male was treated with DMSO and used to generate all lanes. D) Phosphorylation of PKA substrates over time following sperm stimulation with NH_4_Cl, expressed as a fold increase in the intensity of bands from Western blots for phosphorylated PKA substrates. Each point represents the average fold increase for sperm from three unique male genotypes, which were assessed after incubation with DMSO (negative control), the sAC inhibitor KH7, or the PKA inhibitor H-89 for a total of N = 3 assays per point (N = 54 assays in total). Inset identifies time and the interaction between time and treatment (Tr) as significant linear model terms (Type III ANOVA; ** = *p* < 0.01, *** = *p* < 0.001), and bars represent the standard error of the mean.

### Sperm motility under various conditions

Sperm in sea water (SW) exhibited no change in motility after stimulation with 20 mM NH_4_Cl (Tukey’s HSD; *p* = 0.284). For sperm in sodium-free SW (NaFSW), an average of 2 ± 2% of sperm were motile before stimulation, which increased significantly to 69 ± 5% following stimulation with 20 mM NH_4_Cl (*p* < 0.001). A significant increase in motility was also observed for sperm in NaFSW treated with DMSO following stimulation (from 2 ± 2% to 71 ± 6%; *p* < 0.001). In contrast, sperm in NaFSW incubated with KH7 did not show a significant increase in motility after stimulation (from 7 ± 3% to 18 ± 2%; *p* = 0.421). Following incubation with the PKA inhibitor H-89, sperm in NaFSW displayed a significant change in motility after stimulation (from 2 ± 2% to 33 ± 5%; *p* < 0.001). However, the percentage of motile sperm following stimulation in the DMSO treatment was significantly higher than that for the KH7 or H-89 treatments (*p* < 0.001). All sperm motility data are shown in Fig. 5 (N = 9 movies per treatment).

**Figure 5.**
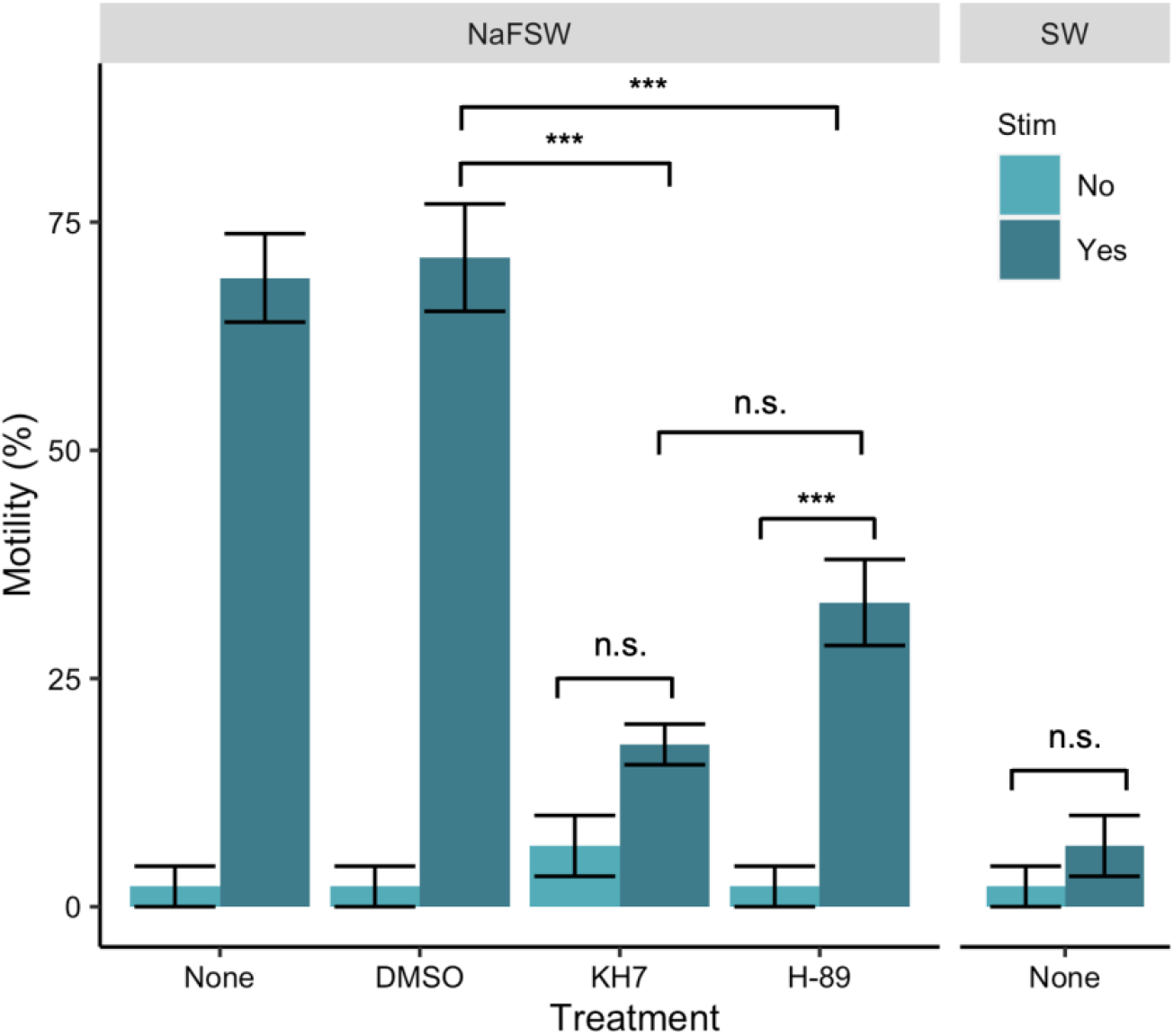
Motility of *Astrangia poculata* sperm under various treatment conditions. Average percent motility observed in movies of sperm swimming in either sodium-free seawater (NaFSW) or regular seawater (SW), and in the presence or absence of 20 mM NH_4_Cl stimulation (“stim”). Sperm were either untreated (“none”) or treated with DMSO (carrier control), the sAC inhibitor KH7, or the PKA inhibitor H-89 prior to recording. Each bar represents an average of three movies (technical replicates) recorded for each of three distinct male colonies, resulting in N = 9 movies per bar or 90 in total. Horizontal lines depict significance of pairwise comparisons (Tukey’s HSD; n.s. = not significant, *** = *p* < 0.001), while vertical error bars indicate the standard error of the mean.

### Sperm motility pathway expression in *A. poculata* proteome

Queries of the *A. poculata* adult proteome resulted in identification of homologs for all proteins constituting the sperm motility pathway in sea urchins and other invertebrates (guanylyl cyclase A, the potassium-selective, cyclic nucleotide gated channel, the hyperpolarization-activated, cyclic nucleotide-gated channel, CatSper α1–4, and the sperm-specific Na^+^/H^+^ exchanger). Each of these homologs was the only significant (E-value < 10^-5^) match for each protein (Table S1). A summary of the function of these proteins with the lengths and predicted molecular weights of the identified *A. poculata* homologs are presented in Table 1.

**Table 1.**
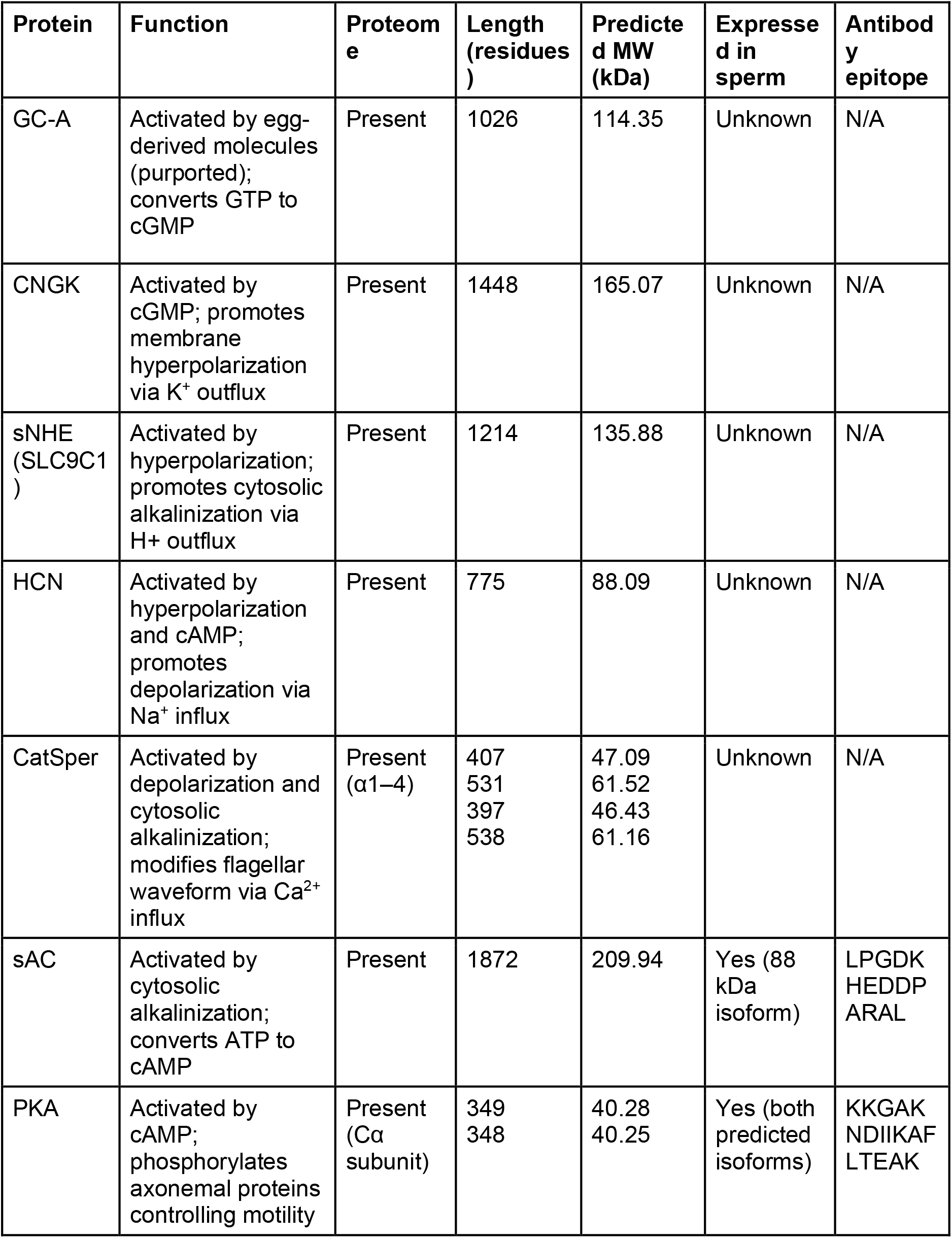
Summary of known sperm motility pathway proteins identified in the *Astrangia poculata* proteome.

### Cnidarian sAC protein phylogeny

A multiple sequence alignment (MSA) of cnidarian full-length sAC protein sequences generated via seeded guide trees and hidden Markov model profile-profile techniques showed high levels of conservation of residues across 12 of 13 representative cnidarian species, which were chosen to represent four major cnidarian clades (octocorals, sea anemones, robust corals, and complex corals). The soft coral *Xenia* sp. expressed a distinct sAC homolog 1100–1700 amino acids shorter than the other species (Fig. S2). The corals *Acropora digitifera* and *Montipora capitata* expressed sAC homologs with extended C-terminal domains 100–300 amino acids longer than the other species, while the corals *Stylophora pistillata* and *Orbicella faveolata* expressed homologs with C-terminal domains 100–500 amino acids shorter than the other corals (Fig. S2). The sea anemone *Nematostella vectensis* displayed a small insertion around amino acids 1000– 1187 not shared by other species in its sAC homolog, as did *S. pistillata* (around amino acids 1617–1878; Fig. S2). The sAC homologues for 12 of the 13 species (again, barring only *Xenia* sp.) contained at least one predicted adenylate/guanylate cyclase catalytic domain (Table 2). The sAC homologue expressed by *Xenia* sp. contained one predicted domain, a tetraricopeptide (TRP) repeat, which was also present in the sAC homologs expressed by *Actinia tenebrosa, Acropora digitifera*, and *Acropora millepora* (Table 2). The sAC homologs for all species except *Xenia* sp. and *O. faveolata* had AAA ATPase domains (Table 2).

**Table 2.**
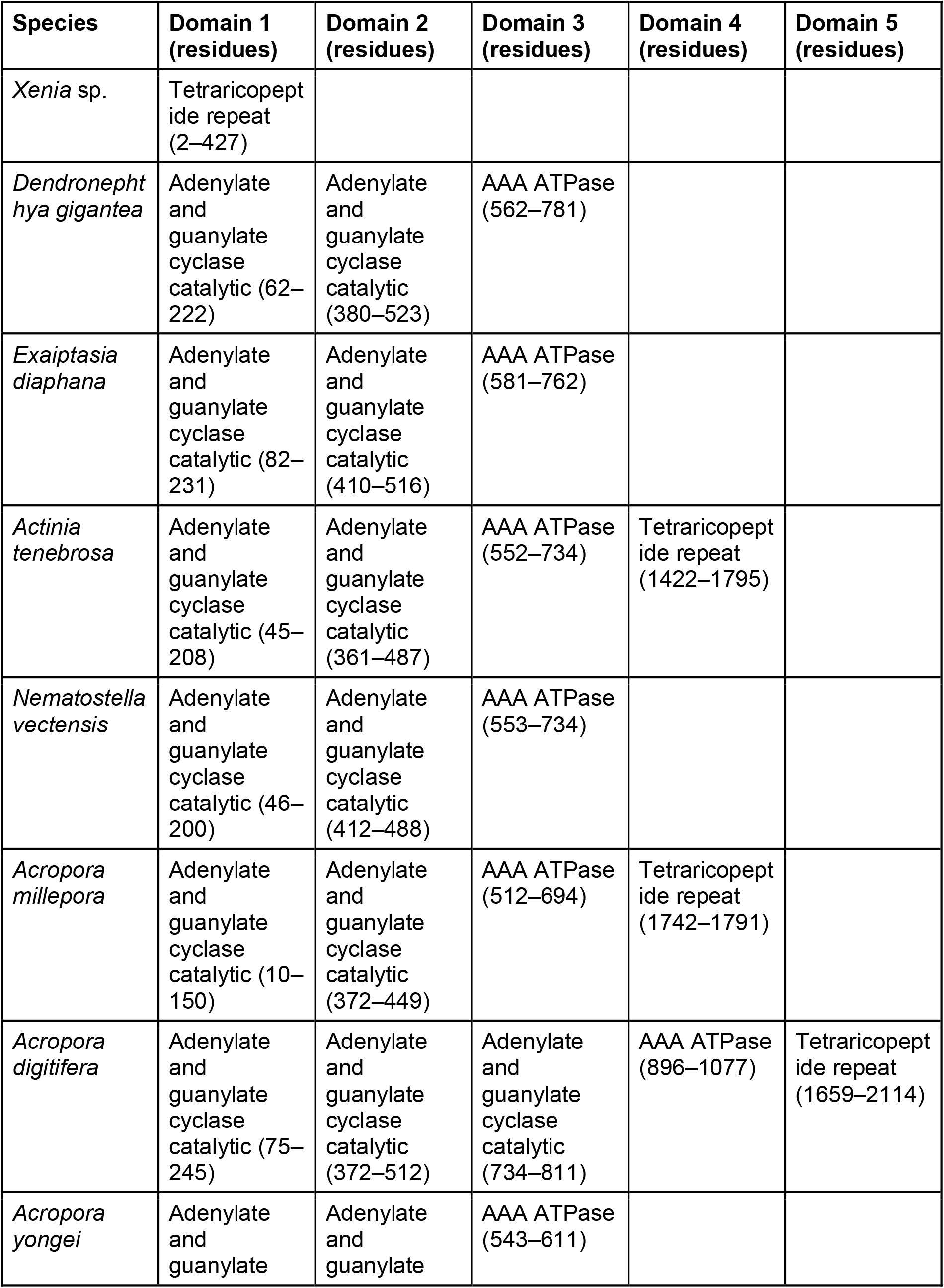
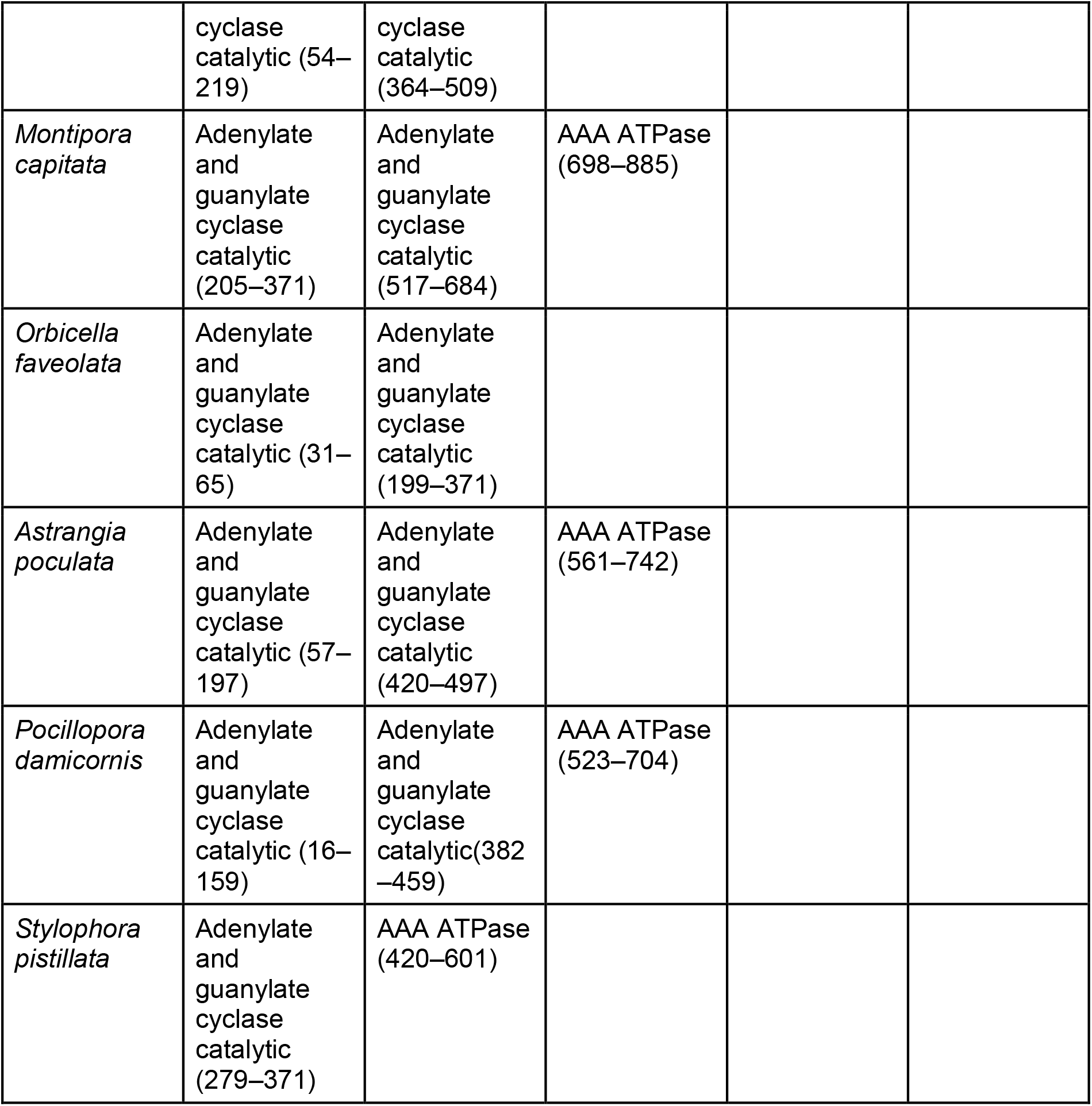
Summary of domain prediction results for cnidarian sAC homologs.

A phylogenetic tree built via neighbor joining algorithms (without distance correction) from the sAC homolog sequences (Fig. 6) indicated that the 13 species clustered into four major groups, which were in line with their accepted phylogenetic relationships, namely octocorals (*Xenia* sp. and *Dendronephthya gigantea*), sea anemones (*Exaiptasia diaphana*, *A. tenebrosa*, and *N. vectensis*), complex corals (*A. millepora*, *A. digitifera*, *A. yongei*, and *M. capitata*), and robust corals (*O. faveolata, A. poculata, Pocillopora damicornis*, and *S. pistillata*). Additionally, accepted phylogenetic relationships (Rodríguez et al., 2014; Zapata et al., 2015) were maintained within each group (e.g. closest grouping of the three *Acropora* species within the complex corals with *M. capitata* as an outgroup), but not between groups, as the robust corals here formed an outgroup to the other three clades (Fig. 6). Furthermore, the stony corals (i.e., robust and complex corals) formed a paraphyletic group on the tree of sAC sequences. Finally, the gonochoric cnidarians *Xenia* sp. (Benayahu, 1991), *Dendronephthya gigantea* (Hwang and Song, 2012), *Exaiptasia diaphana* (Armoza-Zvuloni et al., 2014), *Actinia tenebrosa* (Veale and Lavery, 2012), *Nematostella vectensis*, and *Astrangia poculata* were present in different groups across the tree, as was the case for the hermaphroditic species *Acropora millepora* (Miller et al., 2003), *Acropora digitifera* (Morita et al., 2006), *Acropora yongei* (Zayasu and Suzuki, 2019), *Montipora capitata* (Speer et al., 2021), *Orbicella faveolata* (Steiner, 1991), *Pocillopora damicornis* (Steiner and Cortés, 1996), and *Stylophora pistillata* (Douek et al., 2011).

**Figure 6.**
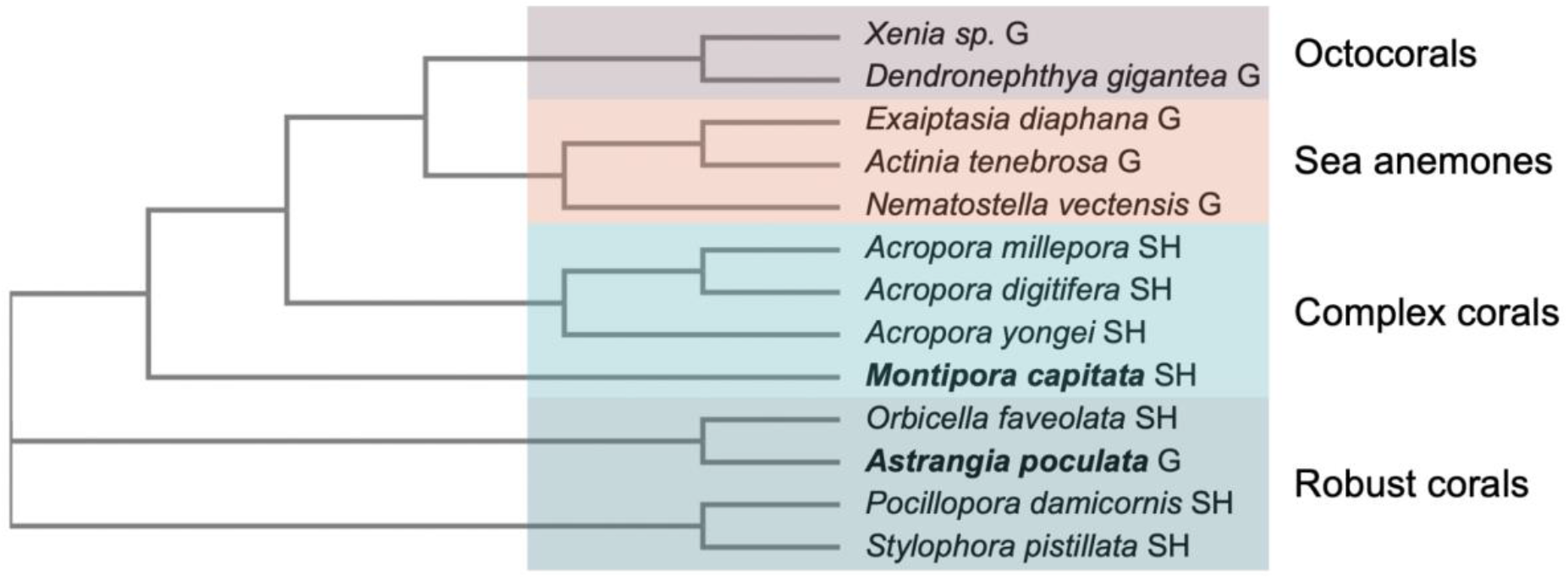
Phylogenetic tree of cnidarian sAC. Phylogenetic tree of 13 cnidarian species built using the amino acid sequences of sAC homologs (predicted sequences or proteome data). Labels next to each name indicate sexual system (G = gonochoric, SH = simultaneous hermaphrodite), and colors depict grouping of recognized clades, which are also labeled. The positions of *Astrangia poculata* and *Montipora capitata* are highlighted by bold text.

## Discussion

In this study, we describe the molecular mechanisms controlling the initiation of sperm motility in the gonochoric coral *Astrangia poculata*, which we found were conserved between this species and other broadcast spawning marine invertebrates (Esposito et al., 2020; Morisawa and Yoshida, 2005; Morita et al., 2009b; Nishigaki et al., 2014; Speer et al., 2021). In addition, we found for the first time in any invertebrate that inhibition of the enzyme soluble adenylyl cyclase (sAC) abolishes the production of cyclic adenosine monophosphate (cAMP) in response of sperm to cytosolic alkalinization, resulting in a lack of downstream sperm motility. We also found that inhibition of sAC reduces protein kinase A (PKA) activity, and that PKA activity is necessary for activation motility in response to cytosolic alkalinization, demonstrating for the first time in any cnidarian that PKA is downstream of sAC and is necessary for sperm motility. Additionally, we present updated ultrastructures of *A. poculata* sperm, which clarify the number of mitochondria and presence of pericentriolar processes. Finally, we queried published cnidarian proteomes and found that sAC, the central signaling node in the sperm motility pathway, demonstrates structural and functional conservation across a diversity of cnidarian species.

### Sperm from *A. poculata* displayed a morphology typical of gonochoric anthozoans

We examined sperm spawned by male *A. poculata* colonies via transmission electron microscopy (TEM), and provide novel insights into ultrastructural features compared to the only prior ultrastructural analysis of sperm from this species published over 40 years ago (Szmant-Froelich et al., 1980). For example, we found that sperm from *A. poculata* contained 2–4 mitochondria, whereas previous work suggested the presence of only a single mitochondrion in sperm from this species (Szmant-Froelich et al., 1980). Furthermore, pericentriolar processes could be seen in our micrographs connecting the central microtubule pair to the outer microtubule doublets in the flagellar axoneme of *A. poculata* sperm, which was not previously reported. Pericentriolar processes are also observed in sperm from other gonochoric anthozoans, and may function to provide structural support to the flagellum (Hinsch, 1974). At a broader level, *A. poculata* sperm displayed all of the ultrastructure features that are characteristic of sperm from gonochoric anthozoans (Steiner, 1991; Steiner, 1993), including: 1) a conically shaped nucleus with a region of low density at the anterior tip (the latter of which may serve as an acrosome); 2) partial fusion of mitochondria resulting in differences in mitochondrial number between sperm cells; 3) the presence of dense perinuclear vesicles; and 4) a dense lipid body planar with the mitochondria. Thus, *A. poculata* sperm display a distinct morphological “type” shared with sperm from other gonochoric but not hermaphroditic corals, the latter of which have rounded nuclei and discrete mitochondria, but lack lipid bodies and perinuclear vesicles (Steiner and Cortés, 1996). As different sperm morphologies correlate with molecular and functional differences both within and outside of phylum Cnidaria (Amaral et al., 2022; Darszon et al., 2020; Steiner, 1991), these results call into question whether cnidarians with different sexual systems utilize the same sperm motility signaling pathway.

### The initiation of *A. poculata* sperm motility is controlled by the sAC-cAMP-PKA pathway

Given their morphological and molecular dissimilarities, we hypothesized that the molecular mechanisms underlying sperm motility would differ between hermaphroditic and gonochoric corals. This hypothesis was surprisingly not supported, as we found that the initiation of sperm motility in *A. poculata* was controlled by a cell signaling pathway involving sAC and PKA, which also controls the onset of sperm motility in the hermaphroditic coral *Montipora capitata* (Speer et al., 2021). We found that, in sperm from *A. poculata*, a single isoform of sAC approximately 88 kDa in weight was expressed in the sperm head as well as throughout the flagellum — a localization that supports sAC’s purported role in regulating flagellar beating that was also observed in *M. capitata* (Speer et al., 2021). Interestingly, the molecular weight of this protein did not match that of the predicted full-length sAC sequence in the *A. poculata* proteome (209.94 kDa), likely due to alternative splicing or post-translational modifications, which may be sperm-specific. Alternative splicing of sAC has been observed in other species including cnidarians and humans (Tresguerres et al., 2014; Xie and Conti, 2004), and the functions of different sAC isoforms could be investigated in future studies.

Next, we found that sperm displayed a burst in cAMP production within six seconds following induced cytosolic alkalinization. Importantly, we showed for the first time in any invertebrate species that this cAMP spike was absent when sperm were pre-treated with the sAC inhibitor KH7, confirming that sAC activity is necessary for this response, as opposed to transmembrane adenylyl cyclases (tmACs). In addition, treatment with KH7 resulted in a significant decrease in motility activation compared to the DMSO control, confirming that sAC activity is necessary for motility in this species. PKA is typically activated downstream of sAC in other taxa (Gancedo, 2013), and here we found that sperm expressed two isoforms of PKA Cα and displayed a rapid increase in PKA substrate phosphorylation following cytosolic alkalinization. PKA activity appeared to peak after the burst in cAMP production, suggesting that PKA might be activated downstream of sAC in this species. In support of this hypothesis, we found that PKA activity was negatively affected by the presence of the sAC inhibitor KH7, providing the first evidence, to our knowledge, that PKA activation occurs downstream of sAC in a cnidarian. Importantly, both PKA activity and sperm motility significantly decreased following treatment with the PKA inhibitor H-89, confirming for the first time that PKA activity is necessary for coral sperm motility, which has been observed in sea urchin sperm (Morisawa and Yoshida, 2005). Surprisingly, sperm pre-treated with KH7 did not show a significant change in motility following induced cytosolic alkalinization, though this was observed for sperm pre-treated with H-89, suggesting that sAC is particularly important for the initiation of motility, while there may be compensatory pathways for activating motility downstream of sAC when only PKA is inhibited. Together, these results demonstrate that induction of the sAC-cAMP-PKA signaling pathway controls the initiation of sperm motility in *A. poculata*.

### The *A. poculata* proteome contains homologs to the entire sea urchin sperm motility pathway

In sea urchins and *M. capitata*, sAC and PKA form part of a larger sperm motility pathway with several transmembrane proteins and other components (Darszon et al., 2020; Speer et al., 2021). We identified homologs of each of the known sperm motility pathway proteins — guanylyl cyclase A (GC-A), the potassium-selective, cyclic nucleotide gated channel (CNGK), the hyperpolarization-activated, cyclic nucleotide-gated channel (HCN), CatSper α1–4, and the sperm-specific Na^+^/H^+^ exchanger (sNHE) — expressed in the proteome of *A. poculata*. While it seems surprising that the sperm-specific CatSper and sNHE proteins were identified in this analysis, the proteome we queried was created using whole adult tissue from several animals, and would likely have included developing spermatocytes (Guiglielmoni, 2021). Thus, we suggest that the entire sperm motility pathway operating in sea urchins and *M. capitata* is also present in *A. poculata*, indicating a surprising level of conservation across species with divergent sexual systems, as well as across phyla. This result has important evolutionary implications, as it suggests that the role of the sAC-cAMP-PKA pathway in controlling sperm motility may have arisen prior to the divergence of early-branching metazoan phyla (e.g. Echinodermata and Cnidaria). However, conservation of this pathway does not preclude differential signaling mechanisms between hermaphroditic and gonochoric species to initiate motility, and more work is needed to characterize the molecules that regulate the initiation of this pathway (e.g. types of ligands and receptors involved) and potential differences in sperm behavior or performance (e.g. longevity, swimming pattern, etc.).

### Cnidarian sAC homologs display broad structural and functional conservation

To further investigate the conservation of the sperm motility pathway across both gonochoric and hermaphroditic cnidarian species, we queried published genome, transcriptome, and proteome datasets for the presence of homologs to sAC, the central signaling node in this pathway, in 13 cnidarian species representing four major clades (octocorals, sea anemones, complex corals, and robust corals). Multiple sequence alignment (MSA) of sAC homologs from these species showed a high degree of amino acid sequence similarity, which correlated with conservation of functional domains including adenylate cyclase and ATPase domains, suggesting that sAC-cAMP-PKA signaling is a mechanism shared by nearly all of these species that may be broadly involved in regulating sperm motility (as we show here for *A. poculata*). Interestingly, a phylogenetic tree of sAC amino acid sequences from these species showed clustering that was neither correlated with sexual system nor accepted phylogenetic relationships (e.g. lack of monophyly for the stony corals), demonstrating a possibly ancient origin for sAC in this phylum, which supports a conserved role for sAC in foundational biological processes such as sexual reproduction. Given that the sAC-cAMP-PKA pathway is pH-sensitive and requires alkalinization of the sperm cytosol for activation, this result also has important implications for broadcast spawning marine invertebrates in the face of anthropogenic ocean acidification (OA), which is already known to alter sperm motility in many species (Byrne et al., 2010; Esposito et al., 2020; Hudson and Sewell, 2022; Morita et al., 2010; Nakamura and Morita, 2012). Future work should investigate the role of sAC-cAMP-PKA signaling in sperm from a broader diversity of species to clarify the mechanisms by which OA might impact reproduction across invertebrate phyla in future seas.

### The molecular sperm motility pathway is conserved across coral sexual systems

Overall, our study demonstrates that the molecular mechanisms underlying sperm motility in the hermaphroditic coral *M. capitata* also operate in the gonochoric coral *A. poculata*, suggesting an unexpected level of conservation of this pathway across sexual systems. We also uncovered broad structural and functional conservation of sAC, the central signaling node in the sperm motility pathway, across a diversity of both gonochoric and hermaphroditic cnidarian species. These results set the groundwork for future research investigating the role of sAC-cAMP-PKA signaling in sperm, while also providing insight into the evolution of the molecular mechanisms underlying sperm motility in an early-branching metazoan clade. Indeed, changes in sperm pHi and concentrations of cyclic nucleotides are associated with sperm motility in hydrozoans, ascidians, sea cucumbers, starfish, sea anemones, corals, and even humans (Hirohashi and Yanagimachi, 2018; Kaupp et al., 2006; Morisawa and Yoshida, 2005; Morita et al., 2006; Morita et al., 2009a; Morita et al., 2009b; Morita et al., 2010; Reuven et al., 2021), suggesting that sperm sAC-cAMP-PKA signaling may be widespread in metazoans. Finally, our work has important conservation implications for corals, which are facing the threat of global extinction due to anthropogenic climate change (van Woesik et al., 2022).

## Materials and methods

### Coral collection and spawning

Adult *Astrangia poculata* (Ellis & Solander, 1786) colonies (Fig. 1A) were collected from Narragansett Bay, RI, USA (41.49231, −71.41883) in early August 2021 and 2022. Corals were transported in a bucket filled with sea water (SW) to the University of Rhode Island Graduate School of Oceanography, where they were cleaned gently by scrubbing any exposed skeleton with a toothbrush to get rid of algae and other attached organisms. After cleaning, corals were put in 24 L plastic tubs (Rubbermaid) filled with SW from Narragansett Bay and equipped with 300 W aquarium heaters (Aqueon). Corals were induced to spawn through temperature ramps (increasing ~22°C–31°C over 30 minutes) and physical touch. When a male coral began to spawn (Fig. 1B), the colony was moved to a small glass bowl (Pyrex) containing 50 mL SW and left unperturbed until spawning activity ceased (Fig. 1C). The colony was then removed and the SW containing live sperm (henceforth referred to as “sperm water”) was passed through a 100 μm cell strainer into a 50 mL conical tube (Thermo Fisher Scientific). This process was repeated for each spawning male, and sperm water was used for downstream assays within four hours after spawning, at which point sperm were confirmed to be alive through visual inspection under a microscope (Nikon Eclipse Ni-U). Whole colonies were imaged before, during, and after spawning using an Apple iPhone 8 (Fig. 1A–C).

### Transmission electron microscopy

Sperm were concentrated by pipetting 2 mL sperm water into a microcentrifuge tube (Eppendorf), followed by centrifugation at 1500 x g for 5 minutes. The supernatant was removed, additional sperm water was added atop the sperm pellet, and centrifugation was repeated. This process was repeated until the sperm pellet was approximately 100 μL, at which point the pellet was fixed with 2.5% (v/v) glutaraldehyde and 2.0% (v/v) paraformaldehyde in 0.1 M sodium cacodylate buffer pH 7.4 (Thermo Fisher Scientific) overnight at 4°C. After subsequent buffer washes, the sample was post-fixed in 2.0% (w/v) osmium tetroxide with 1.5% (w/v) K_3_Fe(CN)_6_ for 1 hour at room temperature, and rinsed in deionized (DI) H_2_O prior to *en bloc* staining with 2% (w/v) uranyl acetate (Thermo Fisher Scientific). After dehydration through a graded ethanol series, the sample was infiltrated and embedded in EMbed-812 (Electron Microscopy Sciences). Thin sections were stained with uranyl acetate and SATO lead, then examined with a JEM-1010 electron microscope (JEOL) fitted with a Hamamatsu digital camera and AMT Advantage NanoSprint500 software.

### Western blotting for sAC and PKA

Sperm pellets were collected from 5–6 different males, individually concentrated in 1.5 mL tubes as previously described, and flash frozen in liquid nitrogen. Samples were then stored at −80°C until analysis. Pellets from six (Aug. 2021) or five (Aug. 2022) male colonies were used for Western blotting according to a standard procedure as described here. Briefly, pellets were lysed in 1x tris-NaCl-EDTA (TNE) lysis buffer, manually dissociated, and sonicated (Diagenode UCD-200) for five minutes with a 30 seconds on/1 minute off cycle to extract proteins, the concentration of which was determined using a standard Bradford assay (Thermo Fisher Scientific). Then, extracted proteins were combined with Laemmli sample buffer (Bio-Rad), denatured at 70°C for 15 minutes, and loaded at 1.08 μg (Aug. 2021) or 1.3 μg (Aug. 2022) protein per well into a 4– 12% tris-glycine gel (Thermo Fisher Scientific). Electrophoresis was performed for 30 minutes at 60 V followed by an hour at 160 V; proteins were transferred to a polyvinylidene fluoride membrane (PVDF; Bio-Rad) under 100 V for 100 minutes at 4°C.

Following transfer, the membrane was blocked for 45 minutes in blocking buffer (5% w/v bovine serum albumin in tris-buffered saline with 0.1% v/v Tween-20; Fisher Thermo Scientific) and incubated with a custom polyclonal primary antibody against coral soluble adenylyl cyclase (sAC; GenScript). This antibody was designed against sAC expressed by the coral *Acropora digitifera* (Barott et al., 2017), and we confirmed that the epitope it targets is 100% conserved in sAC expressed by *A. poculata* but no other proteins from this species. As a nonspecific binding control, a second membrane was incubated with the same primary antibody preabsorbed overnight at 4°C with 300x molar excess of the peptide against which the antibody was designed. The membranes were washed and a secondary antibody (anti-rabbit IgG with horseradish peroxidase; Sigma-Aldrich) was added before final washing, development with pico chemiluminescence reagents (Thermo Fisher Scientific), and imaging on an Amersham Imager 600 (General Electric). Following imaging of sAC bands, the membranes were probed for β-tubulin using monoclonal anti-β-tubulin antibodies (Cell Signaling Technology) and imaged again to serve as a loading control. For blotting of protein kinase A (PKA), the steps described above were repeated with six sperm pellets, and a polyclonal primary antibody against the PKA Cα subunit (Cell Signaling Technology) was used.

### Immunocytofluorescence imaging of sAC in sperm

Immediately following collection, 875 μL sperm water was combined with paraformaldehyde (Thermo Fisher Scientific) at a final concentration of 4% (v/v). Fixed sperm cells were pipetted into square “windows” drawn with a hydrophobic pen on glass microscope slides (Thermo Fisher Scientific), which were incubated at 4°C for 30 minutes and then washed in ice cold phosphate buffered saline (PBS; Sigma-Aldrich). Sperm were permeabilized with 0.1% (v/v) Triton-X in PBS for three minutes followed by blocking with blocking buffer (1% w/v bovine serum albumin with 0.2% w/v keyhole limpet hemocyanin in PBS; Thermo Fisher Scientific) for 30 minutes at 4°C. Next, slides were again washed in PBS and then incubated overnight at 4°C with primary antibodies (anti-sAC and β-tubulin) diluted in blocking buffer. As a nonspecific binding control, each slide contained a window that was incubated with blocking buffer containing no primary antibodies.

Following overnight incubation, slides were washed three times for 10 seconds each in PBS, and then secondary antibodies (anti-rabbit IgG AlexaFluor 488 and anti-mouse IgG AlexaFluor 594; Abcam) diluted in blocking buffer to a concentration of 1:1000 were added to all windows followed by incubation in the dark at room temperature for one hour. Slides were then washed for 10 seconds in PBS and dipped three times for 10 seconds each in clean deionized (DI) H2O prior to addition of mountant (containing DAPI to stain DNA) and glass coverslips to each window. To ensure full drying of the mountant, slides were kept in a dark drawer for 24 hours before storage in the dark at −20°C. Later, slides were allowed to come to room temperature before being imaged on a Leica DMi8 confocal microscope. For each slide, the control window was first used to adjust imaging settings (excitation laser intensity, emission collection ranges, and gain) such that any nonspecific binding of fluorescent markers was eliminated (Fig. S1A); then, windows containing sperm incubated with both primary and secondary antibodies were imaged with the same settings. For imaging of DAPI, samples were excited at 405 nm and emission was collected with a HyD filter in the range 415–512 nm. For imaging of tubulin (AlexaFluor 594), samples were excited at 561 nm and emission was collected with a PMT filter in the range 596–646 nm. For imaging of sAC (AlexaFluor 488), samples were excited at 488 nm and emission was collected with a HyD filter in the range 493–602 nm. Brightfield images were also collected on the same microscope (Fig. S1B).

### Assessment of *in vivo* sAC activity

Following collection of sperm water, pellets from three males were concentrated individually as described above and then resuspended in sodium-free seawater (NaFSW). Suspension in NaFSW disables sperm motility, which can be reactivated by alkalinization of the sperm cytosol via addition of 20 mM NH_4_Cl, which we refer to as “stimulation” (Morita et al., 2009a; Speer et al., 2021). Once resuspended in NaFSW, sperm were incubated with either dimethyl sulfoxide (DMSO; 0.5% v/v) as a carrier control or the sAC inhibitor KH7 (50 μM; Tocris Bioscience) for 30 minutes, then pipetted into a 96-well plate (Thermo Fisher Scientific). For each male colony, replicate wells of sperm treated with DMSO or KH7 were amended with either 20 mM NH_4_Cl (stimulated), or an equivalent volume of NaFSW (unstimulated). Across a six-point time series (time = 0, 0.1, 0.5, 1, 2, and 5 minutes), triplicate wells were lysed with the addition of 0.167 M HCl with 0.01% (v/v) Triton-X. Each plate was covered and stored at −20°C until further analysis. Later, plates were thawed on ice and a standard Bradford assay was performed to determine the protein concentration in each well.

An enzyme-linked immunosorbent assay (ELISA) designed to quantify cAMP was performed using a kit from ArborAssays (K019) according to the manufacturer’s instructions. Briefly, samples were put into a 96-well plate coated with sheep IgG, then a cAMP-peroxidase conjugate was added to each well and the binding reaction was initiated by the addition of a sheep antibody against cAMP to each well. After two hours, the plate was washed and a substrate was added which reacts with the bound cAMP-peroxidase conjugate. Finally, the reaction was stopped and the intensity of each well was measured at 450 nm for comparison to a cAMP standard curve. Data were analyzed in the ArborAssays online portal, which uses a four parameter logistic (4PL) curve fit of the cAMP standards to determine the amount of cAMP (nmol) in each well, which was finally normalized to ng protein in the same well. Normalized cAMP concentrations for each time point, treatment (DMSO or KH7), and stimulation condition were used in statistical analysis and then averaged across the three male colonies for plotting.

### Prediction of PKA Cα 3D structures

Following Western blotting for PKA Cα in *A. poculata* sperm (see above), an available *A. poculata* proteome was queried for expression of PKA Cα. Specifically, a custom protein basic local alignment search tool (BLASTp) database was created from a FASTA file containing the *A. poculata* proteome (Guiglielmoni, 2021). Searching the database using the epitope against which the commercial PKA Cα antibody was designed (KKGAKNDIIKAFLTEAK) yielded two highly similar protein results, which were confirmed to be isoforms of PKA Cα via alignment of their amino acid sequences against human PKA Cα with Clustal Omega (Sievers et al., 2011). Next, predicted molecular weights were calculated for each isoform by adding the individual molecular weights of their constituent amino acids. Finally, 3D structures were generated with AlphaFold (Jumper et al., 2021) and visualized with Mol* (Sehnal et al., 2021).

### Assessment of *in vivo* PKA activity

Sperm from three distinct males were pelleted and resuspended individually in NaFSW as described above. Then, concentrated sperm water from each male was split into three aliquots for incubation with either DMSO (0.5% v/v), the sAC inhibitor KH7 (50 μM), or the PKA inhibitor H-89 (20 μM; Cell Signaling Technologies) for 30 minutes. Following incubation, sperm from each treatment were aliquoted into six 1.5 mL tubes. One tube from each male and treatment was flash frozen after addition of NaFSW as an unstimulated control. Then, 20 mM NH_4_Cl was added to each of the remaining tubes, which were flash frozen in a liquid nitrogen at either 0.1, 0.5, 1, 2, or 5 minutes post-stimulation. For downstream analysis, protein was extracted from each sample, quantified through a standard Bradford assay, and run in gel electrophoresis as described above. Following transfer, PVDF membranes were probed with a commercial primary antibody recognizing only phosphorylated substrates of PKA (Fig. S1C; Abcam). All blots were developed with pico chemiluminescence reagents (Thermo Fisher Scientific), and imaged on an Amersham Imager 600 (General Electric) with auto exposure.

To quantify the activity of PKA following stimulation with NH_4_Cl, each membrane image was analyzed as follows: 1) the blot image was opened in ImageJ (Schneider et al., 2012) and converted to 32-bit grayscale; 2) a rectangular area of interest (AOI) was drawn around the first lane of the blot (time = 0 min or unstimulated) from the top to the bottom of the protein ladder (Fig. S1C); 3) the mean pixel intensity (0–256) for the AOI was measured and recorded; 4) the AOI was moved horizontally to the next lane; and 5) steps 3 and 4 were repeated for each lane. After each image was analyzed, mean pixel intensities were subtracted from 256 such that increased PKA substrate phosphorylation was represented by a larger mean intensity value. Then, the time = 0.1 – 5 min values for each blot were divided by the time = 0 min value for the same blot, resulting in a fold increase in PKA substrate phosphorylation levels for each stimulated time point compared to the unstimulated control.

### Assessment of sperm motility

Sperm were recorded swimming in SW or NaFSW before and immediately after stimulation with 20 mM NH_4_Cl, and either untreated or treated with DMSO (0.05% v/v), KH7 (50 μM), or H-89 (20 μM) for 30 minutes. For each sample, 3 μL of treated sperm were loaded into a well of a Leja microscope slide and then three movies of five seconds each were recorded on an epifluorescence microscope with camera attachment (Nikon Eclipse Ni-U microscope and digital camera system DS-Ri1) under 40x magnification at three positions within the well (front, middle, and rear). This process was repeated for sperm from each of three males. In all movies, most immotile sperm could be seen twitching, indicating viability (Movie S1). To blind observers to the treatment conditions, each movie was renamed with a random string of characters and corresponding treatment information was recorded in a spreadsheet. Three months after renaming, each movie was assigned a score of either 0, 20, 40, 60, 80, or 100% according to the approximate percentage of sperm in the movie displaying progressive, directional motility (Movie S2–3). For analysis, the three movie scores were averaged first within each colony (technical replicates) and then across the three males (biological replicates) to generate a measure of mean percent motility for each treatment condition.

### Investigation of *A. poculata* proteome for sperm motility pathway proteins

The proteome of *A. poculata* (Guiglielmoni, 2021) was investigated via local BLASTp as described above for the presence of proteins making up the canonical sperm motility pathway in other marine invertebrate species, including sea urchins and the coral *Montipora capitata* (Speer et al., 2021). For guanylyl cyclase A (GC-A), the sequence from the coral *Euphyllia ancora* was used as a query (Zhang et al., 2019). For the potassium-selective, cyclic nucleotide gated channel (CNGK) and the hyperpolarization-activated, cyclic nucleotide-gated channel (HCN), sequences from the sea urchins *Arbacia punctulata* and *Strongylocentrotus purpuratus*, respectively, were used as queries (Speer et al., 2021). In the case of the four subunits of CatSper and the sperm-specific Na^+^/H^+^ exchanger (sNHE), sequences from *S. purpuratus* were used with the National Center for Biotechnology Information’s protein BLAST (NCBI BLASTp) database (Johnson et al., 2008) to identify the corresponding protein sequences for the coral *Pocillopora damicornis*, which were then used to query the *A. poculata* database (Speer et al., 2021). When a clear match (E-value = 0.0) was detected, sequences were counted for length in amino acid residues, then the molecular weights were calculated as described above. A summary of query sequences, sources, and E-values for this analysis is provided in Table S1.

### Cnidarian sAC sequence alignment, domain prediction, and phylogenetic tree generation

Cnidarian genome, transcriptome, and proteome data were investigated for the presence of sAC homologs for 13 species representing four major clades (octocorals, sea anemones, complex corals, and robust corals). First, for *A. poculata*, the query sequence LPGDKHEDDPARAL (Barott et al., 2017) was used via local tBLASTn and BLASTp; a single protein was identified, which was confirmed to be sAC via alignment the entire amino acid sequence with that of sAC expressed by the coral *Stylophora pistillata*, which was also included in analysis (Barott et al., 2020). Next, the full-length *A. poculata* sAC sequence was used to query genome and proteome data for the following species: *Xenia* sp. (Lamarck, 1816), *Dendronephthya gigantea* (Verrill, 1864), *Exaiptasia diaphana* (Rapp, 1829), *Actinia tenebrosa* (Farquhar, 1898), *Nematostella vectensis* (Stephenson, 1935), *Acropora millepora* (Ehrenberg, 1834), *A. digitifera* (Dana, 1846), *A. yongei* (Vernon and Wallace, 1984), *Orbicella faveolata* (Ellis and Solander, 1786), and *Pocillopora damicornis* (Linnaeus, 1758), all of which are present on NCBI BLAST. The *A. poculata* sAC sequence was also used via local tBLASTn and BLASTp to identify sAC expressed by *Montipora capitata* (Dana, 1846) from previously published proteome data (Shumaker et al., 2019). Resulting sequences were subjected to domain prediction with InterPro (Blum et al., 2021) and multiple sequence alignment (MSA) generated via seeded guide trees and hidden Markov model profile-profile techniques with Clustal Omega (Sievers et al., 2011), which was also used to generate a phylogenetic tree via neighbor joining. A graphic of the MSA was produced using Jalview version 2.11.2.5 (Waterhouse et al., 2009).

### Data analysis

All statistical analyses and plot generation were performed using RStudio version 2022.7.1.554 (RStudio Team 2022). For statistical analysis of data from *in vivo* sAC activity assays, linear models, generalized linear models, linear mixed effect models, and generalized additive models were created relating the normalized concentrations of cAMP to the factors time, treatment (DMSO or KH7), and stim (presence or absence of 20 mM NH_4_Cl) individually and in each possible additive and interactive combination (e.g. time*treatment or time + treatment). Models were compared for goodness of fit using the Akaike information criterion (AIC) and the best model (lm(cAMP ~ time + treatment*stim)) was subjected to a Type III ANOVA. The same approach was taken for data from *in vivo* PKA activity assays: models were created relating the fold increases in PKA substrate phosphorylation to the factors time and treatment (DMSO, KH7, or H-89), and the best fit model (lm(fold_increase ~ time*treatment)) was subjected to a Type III ANOVA. For data from motility movies, the factors medium (SW or NaFSW), treatment (none, DMSO, KH7, or H-89), and “stim” (presence or absence of NH_4_Cl) were included in models, and the best model (lm(percent_motility ~ treatment*stim)) was subjected to a Type III ANOVA followed by Tukey’s Honest Significant Difference post-hoc tests to investigate significance of pairwise comparisons. Final figures were generated in Microsoft PowerPoint for Mac version 16.66.1. R packages used include: *ggplot2* (Wickham 2016), *plyr* (Wickham 2011), *ggh4x* (van den Brand 2021), *MuMIn* (Bartoń 2022), *lme4* (Bates et al., 2015), *car* (Fox and Weisberg 2019), *emmeans* (Lenth et al. 2022), and *multcomp(view*) (Hothorn et al., 2008).

## Acknowledgements

The authors thank Cassie Raker, Emma Strand, Taylor Lindsay, and Danielle Becker for assistance with coral collection via SCUBA, as well as URI Dive Safety Officer Anya Hanson for diving oversight. The authors also thank Chloé Gilligan and Ariana Huffmyer for assistance with coral spawning. The Electron Microscopy Resource Lab at the University of Pennsylvania (RRID: SCR_022375) provided assistance with the preparation of sperm samples for TEM.

## Data availability

All raw data and R scripts used in data analysis are publicly available on GitHub and Zenodo.

